# A method to estimate contact regions between hands and objects during human multi-digit grasping

**DOI:** 10.1101/2022.09.30.510358

**Authors:** Frieder Hartmann, Guido Maiello, Constantin A. Rothkopf, Roland W. Fleming

## Abstract

In order to grasp an object successfully, we must select appropriate contact regions for our hands on the surface of the object. However, identifying such regions is challenging. Here, we describe a workflow to estimate contact regions from marker-based tracking data. Participants grasp real objects, while we track the 3D position of both the objects and the hand including the fingers’ joints. We first determine joint Euler angles from a selection of tracked markers positioned on the back of the hand. Then, we use state-of-the-art hand mesh reconstruction algorithms to generate a mesh model of the participant’s hand in the current pose and 3D position. Using objects that were either 3D printed, or 3D scanned—and are thus available as both real objects and mesh data—allows us to co-register the hand and object meshes. In turn, this allows us to estimate approximate contact regions by calculating intersections between the hand mesh and the co-registered 3D object mesh. The method may be used to estimate where and how humans grasp objects under a variety of conditions. Therefore, the method could be of interest to researchers studying visual and haptic perception, motor control, human-computer interaction in virtual and augmented reality, and robotics.

**SUMMARY:** When we grasp an object, multiple regions of the fingers and hand typically make contact with the object’s surface. Reconstructing such contact regions is challenging. Here, we present a method for approximately estimating contact regions, by combining marker-based motion capture with existing deep learning-based hand mesh reconstruction.

## INTRODUCTION

The capacity to grasp and manipulate objects is a key ability allowing humans to reshape our environment to our wants and needs. Yet controlling effectively our multi-jointed hands is a challenging task that requires a sophisticated control system. This motor control system is guided by several forms of sensory input, amongst which vision is paramount. Through vision, we identify the objects in our environment as well as estimate their position and physical properties. We can then reach, grasp, and manipulate these objects with ease. Understanding the complex system that links the input at our retinae with the motor commands that control our hands is a key challenge of sensorimotor neuroscience. In order to model, predict, and understand how this system works, we must first be able to study it in detail. This requires high-fidelity measurements of both visual inputs and hand motor outputs.

Past motion tracking technology has imposed a number of limitations on the study of human grasping. For example, systems requiring cables attached to participants’ hands^1, 2^ tend to restrict the range of finger motions, potentially altering the grasping movements or the measurements themselves. Despite such limitations, previous research has been able to identify several factors that influence visually guided grasping. Some of these factors include object shape^3–6^, surface roughness^7–9^, or the orientation of an object relative to the hand^4, 8, 10^. However, to overcome previous technological limitations, a majority of this prior research has employed simple stimuli and highly constrained tasks, thus predominantly focusing on individual factors^3, 4, 6, 7, 10^, two-digit precision grips^3, 4, 6, 9, 11–18^, single objects^19^, or very simple 2D shapes^20, 21^. How previous findings generalize beyond such reduced and artificial lab conditions is unknown. Further, a recent study^22^ has demonstrated, using a haptic glove, that objects can be recognized by how their surface impinges on our hand. This highlights the importance of studying the extended contact regions between our hands and the objects we grasp, not just the contact points between objects and fingertips^23^.

Recent advances in motion capture and 3D hand modeling allow us to move past previous limitations and to study grasping in its full complexity. Passive marker-based motion tracking is now available with millimeter-sized markers that can be attached to the back of participant’s hands to track joint movements^24^. Further, automatic marker identification algorithms for passive marker systems are now sufficiently robust to almost eliminate the need for extensive manual post-processing of marker data^25–27^. Markerless solutions are also reaching impressive levels of performance at tracking animal body parts in videos^28^. These motion tracking methods thus finally allow reliable and non-invasive measurements of complex multi-digit hand movements^24^. Such measurements can inform us about joint kinematics and enable us to estimate contact points between the hand and an object. Additionally, in recent years the computer vision community has been tackling the problem of constructing models of the human hands that can replicate the soft tissue deformations during object grasping and even during self-contact between hand parts^29–32^. Such 3D mesh reconstructions can be derived from different types of data, such as video footage^33, 34^, skeletal joints (derived from marker-based^35^ or markerless tracking^36^) and depth images^37^. One first key advance in this domain was provided by Romero et al.^38^, who derived a parametric hand model (*MANO*) from over 1000 hand scans from 31 subjects in various poses. The model contains parameters for both the pose and shape of the hand, facilitating regression from different sources of data to a full hand reconstruction. The more recent DeepHandMesh^29^ solution builds on this approach by constructing a parametrized model through deep learning and by adding penetration avoidance which more accurately replicates physical interactions between hand parts. By combining such hand mesh reconstructions with 3D tracked object meshes it is thus now possible to estimate contact regions, not just on the surface of objects^39^, but also on the surface of the hand.

Here, we propose a workflow that brings together high-fidelity 3D tracking of objects and hand joints with novel hand mesh reconstruction algorithms. The method yields detailed maps of hand-object contact surfaces. These measurements will assist sensorimotor neuroscientists in extending our theoretical understanding of human visually guided grasping. Further, the method could be useful to researchers in adjacent fields. For example, human factors researchers may use this method to construct better human-machine interface systems in virtual and augmented reality^18^. High-fidelity measurements of human grasping behaviors can also assist roboticists in designing human-inspired robotic grasping systems based on the principles of interactive perception^40–44^. We thus hope that this method will help advance grasping research across neuroscience and engineering fields from sparse descriptions of highly constrained tasks to fuller characterizations of naturalistic grasping behaviors with complex objects and real-world tasks. Our overall approach is outlined in **Figure 1**.

**Figure 1.**
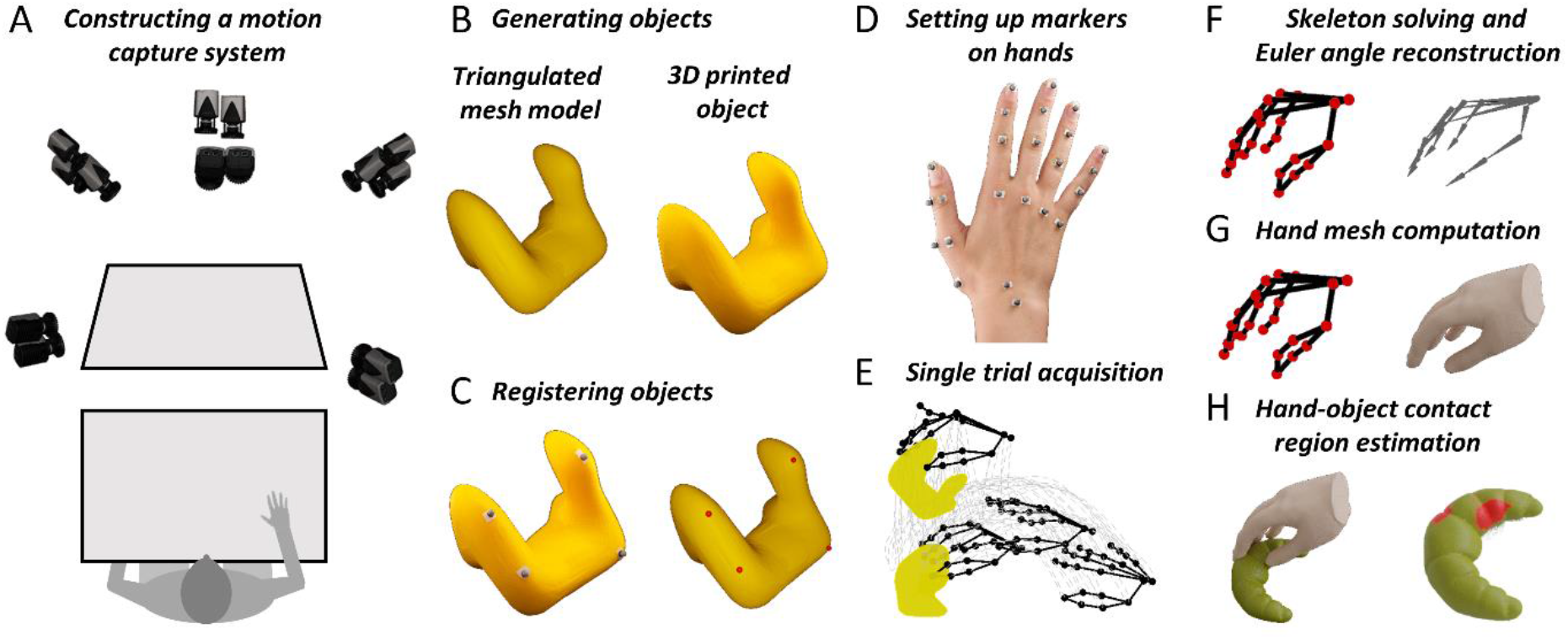
Key steps in the proposed method. **A)** Motion capture cameras image a workbench from multiple angles. **B)** A stimulus object is 3D printed from a triangulated mesh model. **C)** Four spherical reflective markers are glued to the surface of the real object. A semi-automated procedure identifies four corresponding points on the surface of the mesh object. This correspondence allows us to roto-translate the mesh model to the 3D tracked position of the real object. **D)** Reflective markers are attached to different landmarks on the back of a participant’s hand using double - sided tape. **E)** The motion capture system acquires the trajectories in 3D space of the tracked object and hand markers during a single trial. **F)** A participant-specific hand skeleton is constructed using 3D computer graphics software. Skeletal joint poses are then estimated for each frame of each trial in an experiment through inverse kinematics. **G)** Joint poses are input to a modified version of DeepHandMesh^29^, which outputs an estimated 3D hand mesh in the current 3D pose and position. **H)** Finally, we use mesh intersection to compute hand-object contact regions.

## PROTOCOL

Prior to beginning an experiment participants must provide informed consent in accordance with institutional guidelines and the Declaration of Helsinki. All protocols described here have been approved by the local ethics committee of Justus Liebig University Giessen (LEK-FB06).

***Step 1. Install all necessary software***
  **1.1** Download the project repository at: *see Acknowledgments
  **1.2** Install the following software (follow the links for purchase options and instructions):
    **1.2.1** Download and install the Anaconda Python distribution (Anaconda 5.3.1 or later) from https://repo.anaconda.com/archive/ (scripts and functions were generated in Python version 3.7).
    **1.2.1** Within the project repository, run the following command in command window: `conda env create -f environment.yml`
    **1.2.1** MATLAB (scripts and functions were generated in MATLAB version R2018a) https://www.mathworks.com/products/matlab.html
    **1.2.1** Maya https://www.autodesk.com/products/maya/overview
    **1.2.2** QTM Qualisys Track Manager https://www.qualisys.com/software/qualisys-track-manager/
    **1.2.3** Qualisys SDK for Python https://github.com/qualisys/qualisys_python_sdk
    **1.2.3** QTM Connect for Maya https://github.com/qualisys/QTM-Connect-For-Maya
    **1.2.3** Blender 2.92 https://download.blender.org/release/
  **1.3** Download and install the pre-trained DeepHandMesh^29^ instantiation following the instructions provided at: https://github.com/facebookresearch/DeepHandMesh
  **1.4** Place DeepHandMesh in the folder “deephandmesh” of the project repository. Replace the file “main/model.py” with the model.py file contained in the project repository.
***Step 2. Prepare the motion capture system***
  **2.1** Position a workbench within a tracking volume imaged from multiple angles by motion tracking cameras arranged on a frame surrounding the workspace (**Figure 1A**).
  **2.2** Prepare reflective markers by attaching double-sided adhesive tape to the base of each marker.
  **2.3** Calibrate the tracking system following the system-specific calibration routine detailed in the user manual. Before accepting the calibration, export the calibration to a text file.
***Step 3. Create a stimulus object***
  **3.1** Construct a virtual 3D object model in the form of a polygon mesh.
  **3.2** Use a 3D printer to construct a physical replica of the object model. NOTE: In our data repository, we provide example objects in STL and Wavefront OBJ file formats. Objects in STL format are manifold and ready for 3D printing.
***Step 4. Prepare and co-register real and mesh-model versions of the stimulus object***
  **4.1** Attach four non-planar reflective markers to the surface of the real object.
  **4.2** Place the object within the tracking volume.
  **4.3** In the project repository, execute the Python script “Acquire_Object.py”. Follow the instructions provided by the script to perform a 1 second capture of the 3D position of the object markers.
  **4.4** Open MATLAB, navigate to the project repository, and execute the script “Calibrate_Object.m”. Follow the instructions provided by the software routine to identify the marker locations on the object mesh model.
***Step 5. Set up markers on hands***
  **5.1** Attach 24 spherical reflective markers on different landmarks of a participant’s hand using double-sided tape. NOTE: The specific positioning of the markers is demonstrated in **Figure 2**.
    5.1.1 Position markers centrally on top of the respective fingertips, distal interphalangeal joints, proximal interphalangeal joints and metacarpophalangeal joints of the index finger, middle finger, ring finger and small finger.
    5.1.2 For the thumb, position one marker each on the fingertip and basal carpometacarpal joint, and a pair of markers each on the metacarpophalangeal and the interphalangeal joint. These marker pairs need to be displaced in opposite directions perpendicular to the thumb’s main axis and are necessary to estimate the thumb’s orientation.
    5.1.3 Finally, place markers at the center of the wrist and on the scaphotrapeziotrapezoidal joint.
***Step 6. Acquire a single trial***
  **6.1** Ask the participant to place their hand flat on the workbench with the palm facing downwards and to close their eyes.
  **6.2** Place the stimulus object on the workbench in front of the participant.
  **6.3** In the project repository, execute the Python script “Single_Trial_Acquisition.py”. Follow the instructions provided by the script to capture a single trial of a participant grasping the stimulus object. NOTE: The script will produce an auditory cue. This will signal to the participant to open their eyes and execute the grasp. In our demonstrations, the task is to reach and grasp the target object, lift it vertically by approximately 10 cm, set it down, and return the hand to its starting position.
***Step 7. Label markers***
  **7.1** Within QTM, label markers attached to the hand according to the naming convention in Figure 2.
  **7.2** Create an AIM model for the participant’s hand. For successive trials from the same participant, this can be used to automatically identify markers.
***Step 8. Create a personalized skeleton definition for the participant***
  **8.1** Using MAYA and QTM connect for MAYA, load the pre-defined skeleton definition provided in the repository.
  **8.2** Manually fit a hand skeleton to a single frame of motion capture data and export the skeleton definition. NOTE: This step is not strictly necessary, but it is useful to increase the accuracy of the skeleton fits to the marker data. Read the QSolverQuickstartGuide on https://github.com/qualisys/QTM-Connect-For-Maya for more information.
***Step 9. Reconstruct joint skeletal joint poses***
  **9.1** Import the skeleton definition to QTM.
  **9.2** Run the skeleton solver to estimate skeletal joint poses for each frame of each trial in an experiment.
***Step 10. Generate hand mesh reconstructions***
  **10.1** In the project repository, execute the Python script “Hand_Mesh_Reconstruction.py”. Follow the instructions provided by the script to generate, for each frame of the trial a hand mesh reconstructing the current hand pose. NOTE: These mesh reconstructions are generated using a modified version of the open-source and pre-trained hand mesh generation tool, DeepHandMesh^29^.
***Step 11. Generate hand-object contact region estimates***
  **11.1** In the project repository, execute the Python script “Contact_Region_Estimation.py”. Follow the instructions provided by the script to generate hand and object contact region estimates by computing the intersection between hand and object meshes.

**Figure 2.**
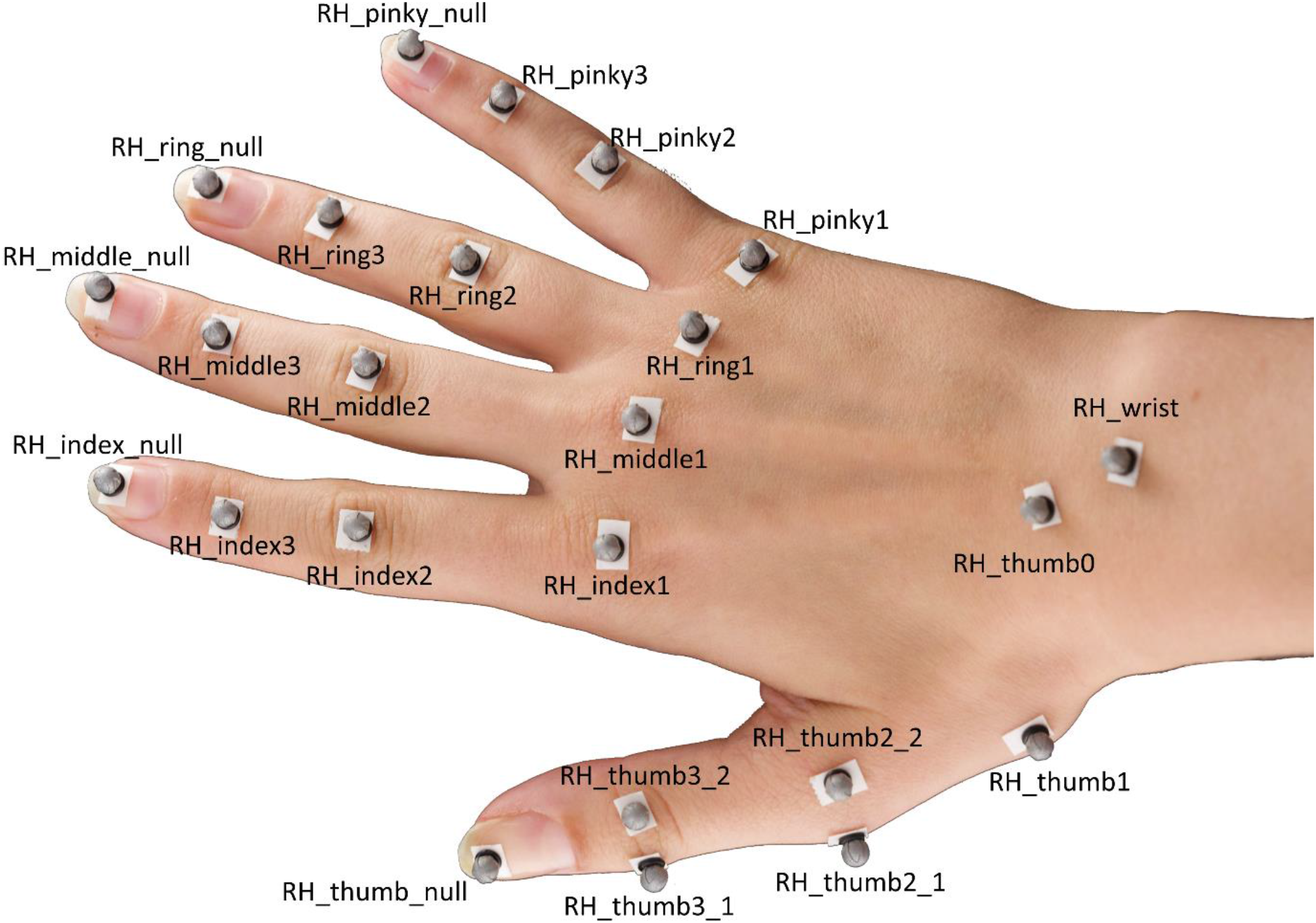
Marker placement on a participant’s hand.

## REPRESENTATIVE RESULTS

The first requirement for the proposed method is a system to accurately track the position of 3D objects and hands. Our specific setup is shown in **Figure 1A**, and uses hardware and software produced by motion capture company Qualisys (Qualisys AB, Sweden). We position a workbench within a tracking volume (100 × 100 × 100 cm) which is imaged from multiple angles by eight tracking cameras (Qualisys Miqus M5) and six video cameras (Qualisys Miqus Video) arranged on a cubical frame surrounding the workspace. The Miqus M5 cameras track the 3D position of reflective markers within the tracking volume at 180 frames per second and with sub-millimeter 3D spatial resolution. We employ 4 mm reflective markers which we attach to objects and hands using skin-friendly double-sided adhesive tape. The 3D marker positions are processed by the Qualisys Track Manager (QTM) software. In the Discussion, we review alternative motion capture systems that could be employed with the proposed method.

To obtain accurate 3D reconstructions of real objects being grasped and manipulated, we propose two options. The first, which is the one we adopt here, is to start from a virtual 3D object model in the form of a polygon mesh. Such 3D models can be constructed using appropriate software (e.g., Blender 3D^45^) and then 3D printed (**Figure 1B**). The second option is to take an existing, real 3D object, and use 3D scanning technology to construct a mesh model replica of the object. Whichever the strategy, the end goal is to obtain both a real 3D object and the corresponding virtual 3D object mesh model. Note that the approach we describe here works only with rigid (i.e., non-deformable) objects.

Once the 3D surface of an object is available as a mesh model, its position must be tracked and co-registered (**Figure 1C**). To do so, we first attach four non-planar reflective markers to the surface of the real object, and place the object within the tracking volume. We then perform a brief capture of the 3D position of the object markers. We use this capture to establish the correspondence between the four markers and four vertices of the object mesh model. This is done using a simple ad-hoc software route written in MATLAB. The program presents a user with a 3D rendering of the virtual object. The user can rotate the rendering in 3D to identify the approximate marker locations. Once the user records these approximate locations, the program constructs 1 cm search spheres centered on these approximate locations. The routine then searches, across every possible combination of four object vertices contained within these spheres, for the best-fitting partial Procrustes alignment (without scale and reflection components) between the four vertices and the positions of the four tracked markers. This procedure yields the four vertices on the object mesh that best correspond to the four tracked object markers. Note that this assumes that the mesh has sufficient resolution that marker locations are well approximated by the nearest vertices (for very low resolution meshes, the nearest vertex could be a poor approximation). Future implementations could include estimating interpolated surface locations between vertices.

Having established correspondence, each time the real object is moved within the tracking volume, the virtual object can be placed in the new position by computing the roto-translation between the tracked markers and the four corresponding mesh vertices (e.g., again using Procrustes analysis). To record the dynamics of the grasp instead, we attach a total of 24 spherical reflective markers on different landmarks of the hand using double-sided tape (**Figure 1D** and **Figure 2**).

At the beginning of a trial (**Figure 1E**), a participant places their hand flat on the workbench with the palm facing downwards and closes their eyes. The experimenter places a target object on the workbench in front of the participant. Next, an auditory cue signals to the participant to open their eyes and execute the grasp. In our demonstrations, the task is to reach and grasp the target object, lift it vertically by approximately 10 cm, set it down, and return the hand to its starting position. A script written in Python 3.7 controls the experiment. On each trial, the script selects and communicates the current condition settings to the experimenter (e.g., object identity and positioning). The script also controls trial timing, including auditory cues and the start and stop of the motion capture recordings.

Limbs are not only characterized by their position in 3D space, but also by their pose. Thus, to obtain a complete 3D reconstruction of a human hand executing a real grasp, we require not only the positions of each joint in 3D space, but also the relative pose (translation and rotation) of each joint with respect to its parent joint (**Figure 1F**). Skeletal joint positions and orientations can be inferred from marker positions using inverse kinematics. To do so, here we employ the skeleton solver provided by the QTM software. For the solver to work, we must first provide a skeleton definition that links the position and orientation of each joint to multiple marker positions. We construct such a skeleton definition and link the skeleton rig to the maker data using the QTM Connect plugin for Maya (Maya, Autodesk, Inc.). We create personalized skeleton definitions for each participant in order to maximize the accuracy of the skeleton fits to the marker data. For each participant, we manually fit a hand skeleton to a single frame of motion capture data. Having obtained a participant-specific skeleton definition, we then run the skeleton solver to estimate skeletal joint poses for each frame of each trial in an experiment.

For each frame of each trial in an experiment, we generate a hand mesh reconstructing the current hand pose using the open-source and pre-trained hand mesh generation tool, DeepHandMesh^28^ (**Figure 1G**). DeepHandMesh is a deep encoder-decoder network that generates personalized hand meshes from images. First, the encoder estimates the pose of a hand within an image, i.e., the joint Euler angles. Then, the estimated hand pose and a personalized ID vector are input to the decoder, which estimates a set of three additive correctives to a rigged template mesh. Finally, the template mesh is deformed according to the estimated hand pose and correctives using Linear Blend Skinning. The first corrective is an ID-dependent skeleton corrective through which the skeletal rig is adjusted to incorporate the person-specific joint positions. The other two correctives are mesh correctives through which the mesh vertices are adjusted to better represent the hand surface of the participant. One of the mesh correctives is an ID-dependent mesh corrective accounting for the surface structure of an individual participant’s hand. The final mesh corrective instead is a pose-dependent vertex corrective which accounts for hand surface deformations due to the current hand pose.

DeepHandMesh is trained using weak supervision using 2D joint key points and scene depth maps. Here, we use only the pre-trained DeepHandMesh decoder to generate hand mesh reconstructions, which we modified in the following ways (**Figure 3**). First, as we do not train the network on our specific participants, we employ the generic ID-dependent mesh corrective provided with the pre-trained model (**Figure 3A**). Further, we derive the ID-dependent skeleton corrective using the QTM skeleton solver as described above (**Figure 3B**). We assume proportional scaling of the hand with skeleton length, and uniformly scale the mesh thickness by a factor derived from the relative scaling of the skeleton, such that the mesh better approximates the participant’s hand size (**Figure 3C**). We input this modified mesh to the decoder, together with the current hand pose (derived from our marker data), and the 3D position and orientation of the wrist. The decoder thus computes the current pose-dependent corrective, applies all correctives and roto-translations, and outputs a 3D hand mesh reconstruction of the current hand pose in the same coordinate frame as the 3D tracked object mesh (**Figure 3D**).

**Figure 3.**
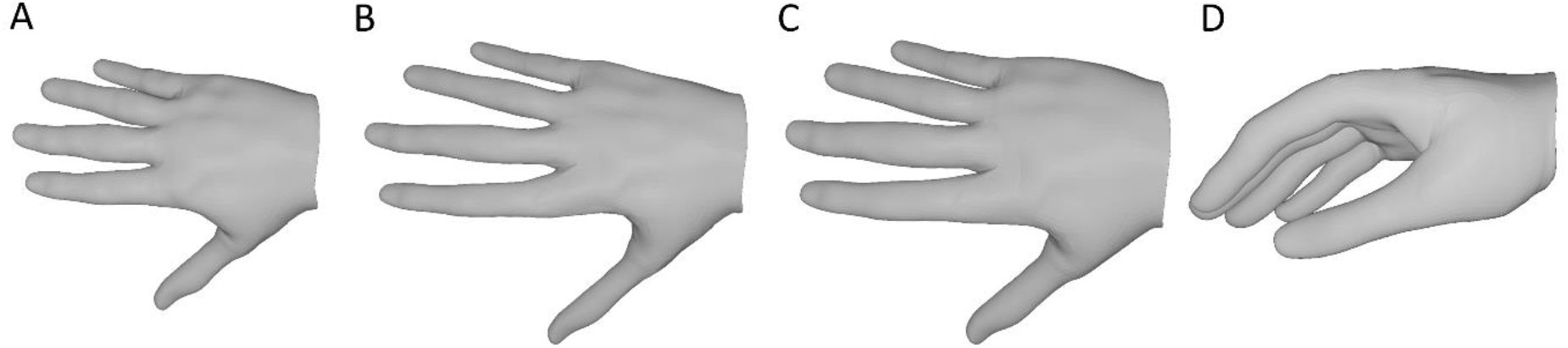
Modifications to the pre-trained DeepHandMesh decoder. **A)** Fixed, generic ID-dependent mesh corrective. **B)** ID-dependent skeleton corrective derived through inverse kinematics in Step 9. **C)** The size of the hand mesh is scaled by the same factor as the skeletal joints. **D)** Final 3D hand mesh reconstruction of the current hand pose.

**Figure 4.**
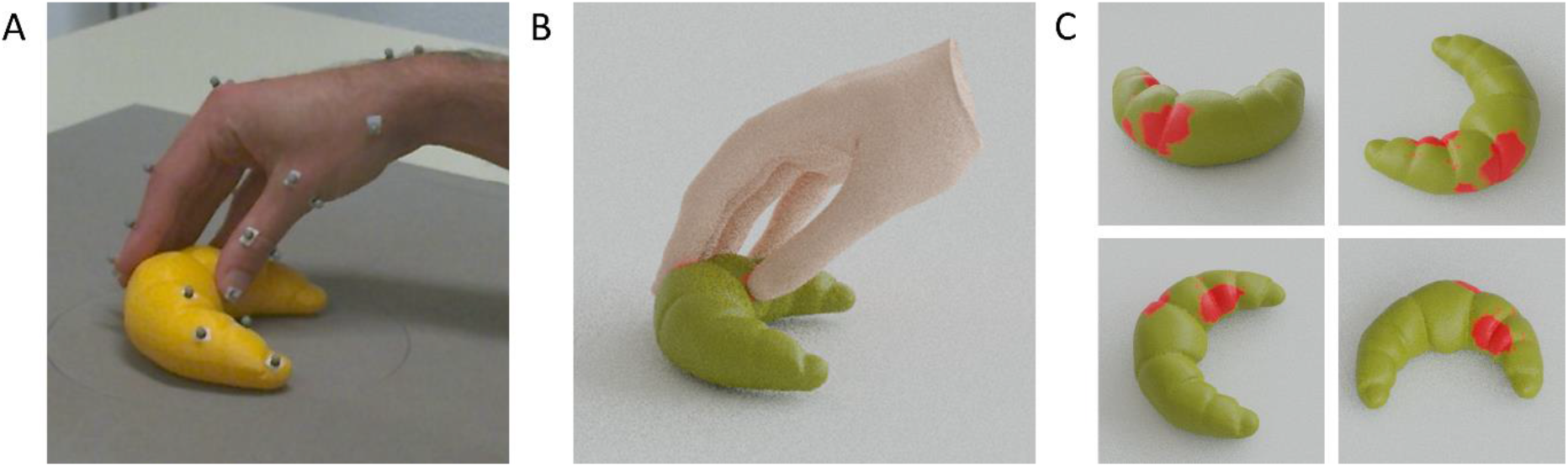
Estimated hand-object contact regions. **A)** Tracked hand and object viewed from one of the tracking cameras during a grasp. **B)** Reconstructed hand mesh and tracked object mesh rendered from the same viewpoint as the tracking camera. **C)** Contact regions on the surface of the object seen from multiple viewpoints.

Having reconstructed 3D mesh models for both a participant’s hand and a grasped object we can then estimate hand-object contact regions by computing the intersection between hand and object meshes (**Figure 1H**). The assumption behind this is that the real hand is deformed by contact with the surface, allowing the skeleton to come closer to the surface than would be possible if the hand were rigid, allowing portions of the hand mesh to pass through the object mesh. As a result, the contact areas can be approximated as the regions of overlap between the two meshes.

Specifically, to compute these regions of overlap we define object mesh vertices that are contained within the 3D volume of the hand mesh as being in contact with the hand. We identify these vertices using a standard raytracing approach^46^. For each vertex of the object mesh, we cast a ray from that vertex to an arbitrary 3D point outside of the hand mesh. We then assess the number of intersections occurring between the cast ray and the triangles composing the hand’s surface. If the number of intersections is odd, the object vertex is contained inside the hand mesh. If the number of intersections is even, then the object vertex is outside the hand mesh. The contact regions on the surface of the object can thus be approximated as the set of triangle faces whose vertices are all contained within the hand mesh. We can apply the same rationale to hand mesh vertices contained in the 3D volume of the object mesh to estimate contact regions on the surface of the hand. Note that more advanced approaches to Boolean mesh operations could also be used^31^.

**Movie 1** shows a video of hand, tracked points, and co-registered mesh, all moving side-by-side during a single grasp to a 3D printed cat figurine. **Figure 4A** instead shows a single frame at the time of hand-object contact from a grasp to a 3D printed croissant, together with the hand/object mesh reconstructions (**Figure 4B**), and the estimated contact regions on the surface of the croissant (**Figure 4C**).

**Movie 1.**
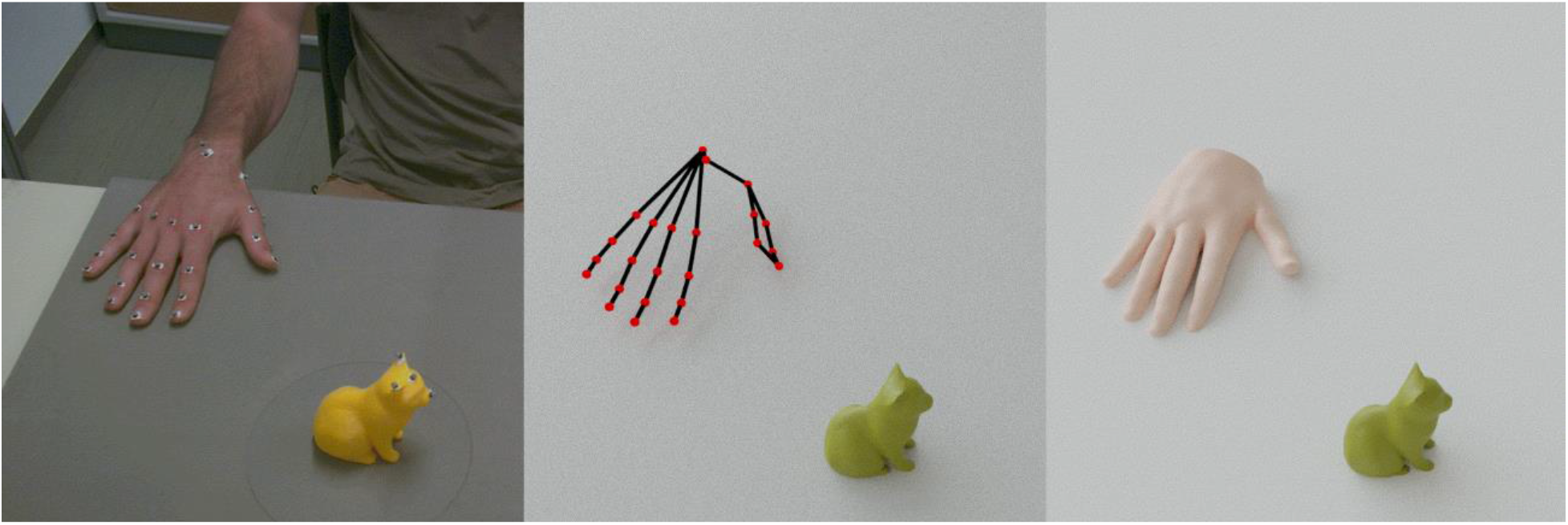
Mesh reconstructions of the hand and object. Gif animation of the hand, tracked markers and the hand and object mesh reconstructions during a single grasp viewed from the same camera viewpoint.

## DISCUSSION

We propose a method that enables estimating contact regions for hand object interactions during multi-digit grasps. Because full tracking of the whole surface of a hand is currently intractable, we propose using a reconstruction of a hand mesh whose pose is determined by sparse keypoints on the hand. To track these sparse keypoints, our solution employs a research-grade motion capture system based on passive marker tracking. Of course, other motion capture systems could also be employed with the proposed method, granted that they yield sufficiently accurate 3D position data. We advise against active marker motion capture systems (such as the popular but discontinued Optotrak Certus by Northern Digital), since these require attaching cables and/or electronic devices to participant hands, which may restrict movements, or at least yield less typical grasps as participants are made more consciously aware of the pose of their hands. Motion tracking gloves using inertial measurement units may be a possibility, even though these systems are known to suffer from drift, may also restrict hand movements, and do not allow for the surface of the hand to come into full and direct contact with object surfaces. Commercial markerless hand tracking solutions (e.g. the Leap Motion^47–49^, now Ultraleap Limited, England) may also be a possibility, although with these systems alone it may not be possible to track object positions. In our opinion, the most promising alternative option to a research-grade motion capture system is given by open source markerless tracking solutions (e.g.^28^). If used with multiple co-registered cameras^50^, such systems could potentially track hand joint positions as well as object positions in 3D, without the need of markers, gloves or cables. These solutions, as well as our marker-based system, may suffer however from data loss issues due to occlusions.

### Limitations and future directions

Because hand reconstructions obtained through our method will not be fully accurate, there are some limitations on the types of experiments for which the method should be used. Deviations of hand mesh reconstructions from ground truth will manifest themselves in deviations of estimated hand/object contact regions. Thus, care should be taken in applying our method to derive absolute measures derived from contact regions. However, even an approximative measure can still be useful in within-subject experiment designs because potential biases of the method will likely affect different experimental conditions within a participant in a similar way. Therefore, we advise only to consider measures such as the differences in contact area between conditions for any statistical analyses as the direction of any effects will correlate with respective ground truth.

While most processing steps from data collection to the final contact region estimation are fully automated, and thus offer important contributions towards a standardized procedure for hand/object contact region estimation, an initial fit of the individualized skeletons to the 3D positions of the tracked markers must still be performed manually to obtain a skeleton definition for each participant. As the number of participants for an experiment increases, so does the number of manual fits which currently is the most time-consuming step in the procedure and requires some familiarity with manual rigging in the Autodesk Maya Software. In the future, we aim to automate this step to avoid human influence on the procedure by adding an automatic skeleton calibration procedure.

Another important limitation of the method is that in its current form, it can only be applied to rigid (non-deformable) objects. In the future, this limitation could be overcome using methods for recording the surface shape of the grasped object as it deforms. Additionally, due to its approximate nature, the method is not currently well suited to very small or thin objects.

In conclusion, by integrating state-of-the-art motion tracking with high-fidelity hand surface modeling we provide a method to estimate hand-object contact regions during grasping and manipulation. In future research, we plan to deploy this method to investigate and model visually guided grasping behavior in humans^16^. We further plan to integrate these tools with eye tracking^47, 51–53^ and virtual/augmented reality systems^54–56^ to investigate visually guided hand and eye movement motor control in real and virtual naturalistic environments^18, 47, 57, 58^. For these reasons, the proposed method could be of interest to researchers studying haptic perception^59^, motor control, and human-computer interaction in virtual and augmented reality. Finally, accurate measurements of human grasping abilities could inform the design of robust robotic systems based on the principles of interactive perception^40–44^, and may have translational applications for upper limb prosthetics.

## ACKNOWLEDGMENTS

Research funded by Deutsche Forschungsgemeinschaft (DFG, German Research Foundation: project No. 222641018-SFB/TRR 135 TP C1 and IRTG-1901 “The Brain in Action”), and by the Research Cluster “The Adaptive Mind” funded by the Excellence Program of the Hessian Ministry of Higher Education, Science, Research, and Art. The authors thank the Qualisys support team, including Mathias Bankay and Jeffrey Thingvold, for assistance in developing our methods. The authors also thank Michaela Jeschke for posing as hand model.

All data and analysis scripts to reproduce the method and the results presented in the manuscript will be made available on a public repository upon acceptance. The repository is currently under construction, but those interested should email us to request the current and most up-to-date version of the repository.

## DISCLOSURES

The authors declare that no competing interests exist.

## Notes

### Competing Interest Statement

The authors have declared no competing interest.

