## Supplementary figures and images for "A method to estimate contact regions between hands and objects during human multi-digit grasping"

### Movie 1

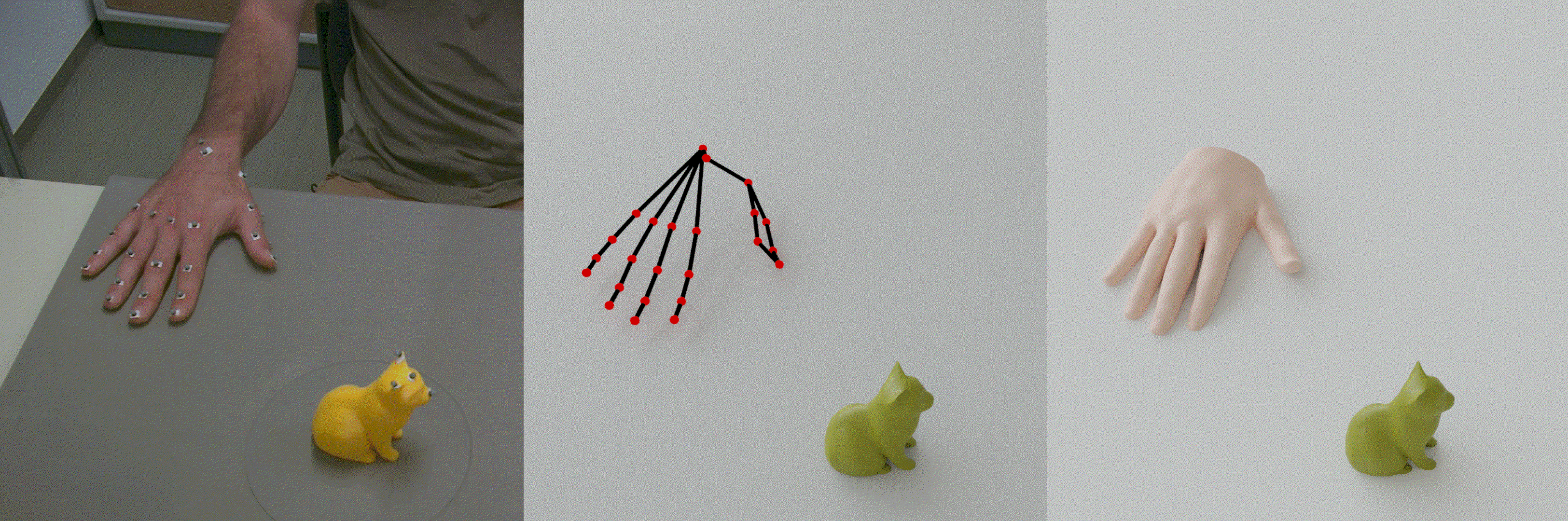
